# High-Resolution X-ray Structure of Gln143Asn Manganese Superoxide Dismutase Captures Multiple Hydrogen Peroxide Binding Sites

**DOI:** 10.1101/2025.07.17.665311

**Authors:** Medhanjali Dasgupta, Katelyn Slobodnik, Erika A. Cone, Jahaun Azadmanesh, Thomas Kroll, Gloria E.O. Borgstahl

## Abstract

Human mitochondrial manganese superoxide dismutase (MnSOD) converts superoxide (O ^●−^) into hydrogen peroxide (H O ) and molecular oxygen (O_2_), serving as a key defense against oxidative damage. Despite extensive studies, the full structural characterization of H_2_O_2_-binding sites in MnSOD remains largely unexplored. Previous H_2_O_2_-soaked MnSOD structures have identified two distinct H_2_O_2_-binding sites: one directly ligated to the catalytic Mn (LIG position) and another at the active site gateway (PEO position) between second-shell residues Tyr34 and His30. In this study, a kinetically impaired Gln143Asn MnSOD variant is used to trap and explore additional H_2_O_2_-binding sites beyond the second-shell solvent gate. In the wild-type enzyme, Gln143 mediates proton transfers with the Mn-bound solvent (WAT1) to drive redox cycling of the metal, necessary for effective O ^●−^ dismutation. Substitution with Asn stalls catalysis because the increased distance from WAT1 disrupts critical proton-coupled electron transfer (PCET) events, and the redox cycling of the active site metal is impaired. This, in turn, stalls the electrostatic cycling of positive charge on the enzyme surface and enhances the likelihood of trapping transient H_2_O_2_-bound states in this variant. Results reveal several H_2_O_2_ molecules leading up to the active site, in addition to the canonical LIG and PEO positions.

**Synopsis:** A high-resolution X-ray structure of a Gln143Asn variant of manganese superoxide dismutase reveals multiple hydrogen peroxide binding sites beyond the canonical LIG and PEO binding positions within the active site. These findings expand the known landscape of product peroxide-bound states in MnSOD.

## 1. Introduction

Human manganese superoxide dismutase (MnSOD) is a mitochondrial matrix metalloenzyme that catalyzes the dismutation of reactive superoxide radicals (O_2_^●−^), generated as by-products of mitochondrial respiration, into molecular oxygen (O_2_) and hydrogen peroxide (H_2_O_2_) (Abreu & Cabelli, 2010). As the only known mitochondrial enzyme capable of converting reactive O_2_^●−^ radicals into diffusible products that can be further metabolized or exported, MnSOD serves as the critical first line of defense against oxidative stress (Bresciani *et al*., 2015, Grujicic & Allen, 2025). Unsurprisingly, dysregulation of MnSOD activity is associated with various human diseases, highlighting the continued need for a detailed structural understanding of its catalytic mechanism.

Human MnSOD physiologically functions as a homotetramer, with each active site containing a Mn ion, which cycles between oxidized (trivalent, Mn^3+^, pink) and reduced (divalent, Mn^2+^, colorless) states to drive two half-reactions to complete the catalytic cycle (Borgstahl *et al*., 1992, Fee & Bull, 1986, Hsu *et al*., 1996).

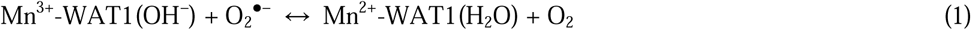

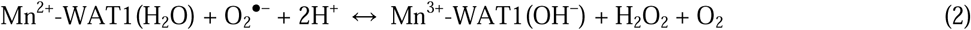

In the first half-reaction (Eq. 1), resting-state Mn^3+^ oxidizes O_2_^●−^ to the first reaction product, O_2_, which diffuses out of the mitochondria or gets reabsorbed into the mitochondrial electron transport chain. During this half reaction, the active site Mn is simultaneously reduced, and its WAT1 ligand receives a proton and becomes a water molecule (Azadmanesh *et al*., 2021). In the second half-reaction (Eq. 2), Mn^2+^ binds a second O_2_^●−^ radical, again protons and electrons are transferred simultaneously to form (H_2_O_2_ and to redox cycle the active site Mn. In this way, both half reactions use proton-coupled electron transfer (PCET) events (Reece & Nocera, 2009), to rapidly convert O_2_^●−^ to O_2_ and H_2_O_2_.

Decades of extensive structural characterization efforts have yielded 39 total human MnSOD structures in the Protein Data Bank (PDB). Out of them, almost all reported H_2_O_2_-soaked MnSOD structures have consistently identified H_2_O_2_ bound to the catalytic Mn, in a near side-on configuration, that replaces the WAT1 solvent (Azadmanesh *et al*., 2025, Azadmanesh *et al*., 2024, Porta *et al*., 2010). This metal-ligated H_2_O_2_ binding site is typically referred to as the LIG position. A recent H_2_O_2_-soaked Trp161Phe MnSOD neutron structure identified a second H_2_O_2_ binding site, named PEO, between the second-shell gateway residues, Tyr34 and His30, at the opening of the active site solvent funnel (Azadmanesh *et al*., 2025). While these structures offer valuable insights into active site H_2_O_2_ binding modes in MnSOD, to date, no electron density for H_2_O_2_ has been observed beyond the PEO binding site, on the solvent-exposed side of the active site funnel. This absence of any observable H_2_O_2_ binding beyond the MnSOD solvent gate, even in severely product-inhibited variants such as Trp161Phe and Tyr34Phe MnSODs, suggests that once HlJOlJ crosses the Tyr34-His30 gateway, it rapidly escapes into bulk solvent.

Further complicating structural efforts, partially occupied or mobile H_2_O_2_ molecules are often indistinguishable from water (H_2_O) in medium-to low-resolution X-ray maps, particularly in the absence of neutron diffraction data that can accurately resolve hydrogen positions. Thus, key questions remain unresolved: (i) What is the fate of H_2_O_2_ following release from the PEO site? (ii) Are there any transient H_2_O_2_ binding interactions occurring beyond the Tyr34-His30 solvent gateway? (iii) What governs H_2_O_2_ diffusion on the solvent-exposed surface of MnSOD? These questions are important because MnSOD catalysis is limited not by the chemical transformation itself, but by diffusion of substrate and product through the ∼10 Å deep, ∼5 Å wide active solvent cavity (Azadmanesh *et al*., 2017).

The goal of this current study is to identify all possible H_2_O_2_ binding sites on the solvent-exposed side of the MnSOD solvent funnel, by using a kinetically slow and catalytically impaired Gln143Asn MnSOD variant (Hsieh *et al*., 1998). Gln143 is a second sphere residue that plays a critical role in the PCET reaction mechanism of native MnSODs (Azadmanesh *et al*., 2021). Azadmanesh and coworkers showed that in the first half-reaction, upon chemical reduction of Mn^3+^ to Mn^2+^, Gln143 donates a proton to the active site metal-bound solvent (WAT1-OH^−^) to generate WAT1-H_2_O (Eq. 1). The WAT1-OH^−^ negative charge is thought to have stabilized the positive charge of Mn^3+^ in the oxidized state. Presumably, in the second half of the reaction (Eq. 2), Gln143 takes the proton back from WAT1-H_2_O, regenerating the WAT1-OH^−^ negative charge simultaneously as Mn^2+^ is oxidized to Mn^3+^. Through this critical and reversible proton transfer event, Gln143 is directly coupled to the redox cycling of the catalytic metal in MnSOD that drives its catalysis (Azadmanesh *et al*., 2021). Substituting Gln143 with Asn preserves the active site architecture, but the hole created disrupts proton transfers with the WAT1 solvent. Consequently, redox cycling of the catalytic Mn is slowed, as evidenced by over 100-fold reductions in the corresponding kinetic rate constants, k_2_ and k_4,_ relative to the wild-type MnSOD (Hsieh *et al*., 1998) (Fig. 1). The kinetic rate constants of Gln143Asn MnSOD have the overall effect of tending to keep the active site Mn reduced. This significantly impairs Gln143Asn MnSOD catalytic turnover with a ∼13-fold decrease in k_CAT_ value relative to wild-type MnSOD (wild-type k_CAT_ = 40 ms^-1^, Gln143Asn k_CAT_ = 0.3 ms^-1^). While this loss of catalytic efficiency compromises the overall function of Gln143Asn MnSOD, it offers a distinct experimental advantage for our study; the slowed turnover rate allows for greater control of experimental design, cryo-trapping, and structural characterization of H_2_O_2_ molecules bound. Our results reveal many H_2_O_2_ molecules beyond the canonical LIG and PEO positions, spanning the space extending from the edge of the Tyr34-His30 solvent gateway to the bulk solvent. These findings expand the structural landscape of how H_2_O_2_ exits the MnSOD active site.

**Figure 1.**
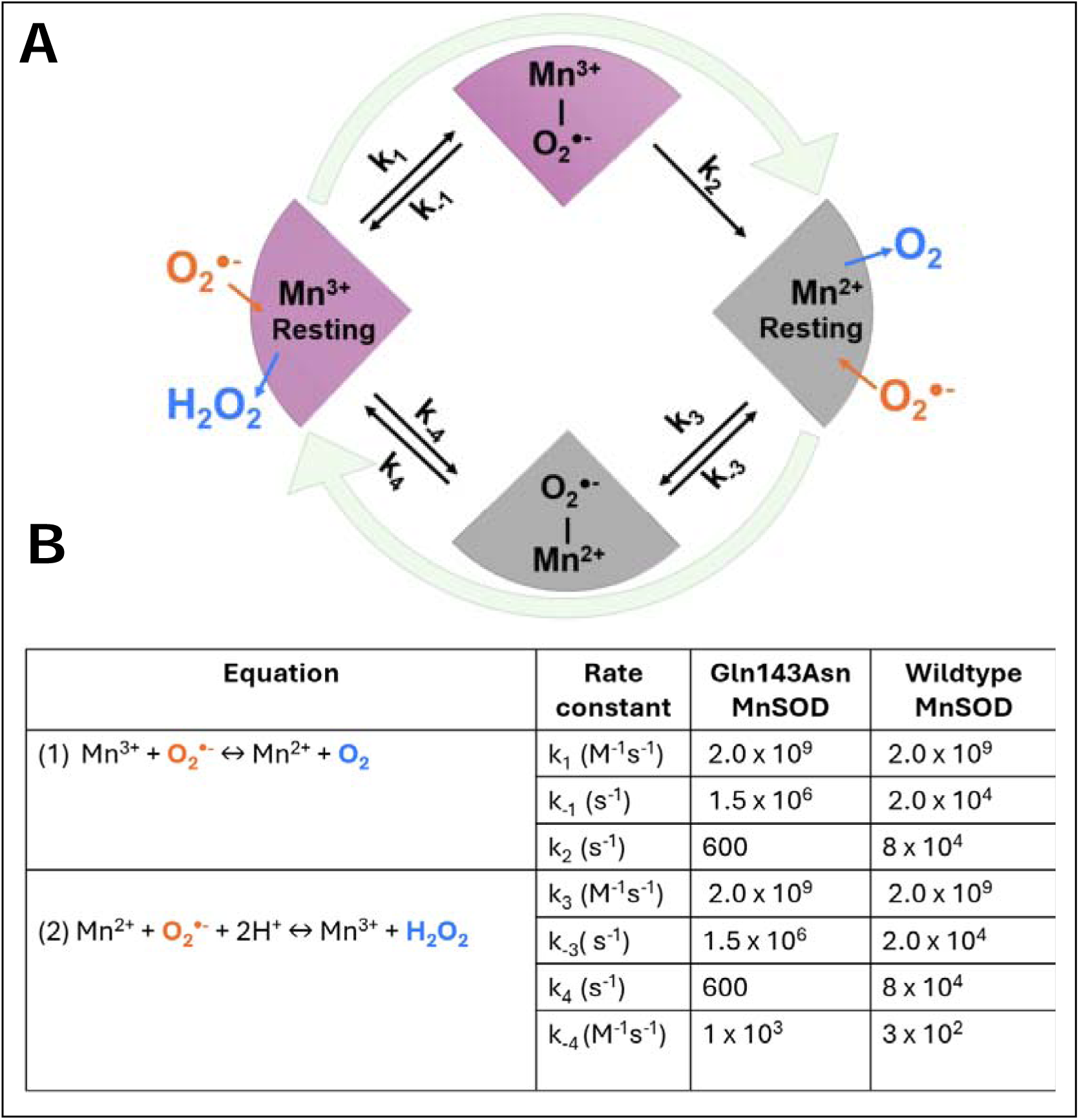
The kinetics of MnSOD catalysis. **(A)** MnSOD reaction scheme, **(B)** Kinetic parameters of human wild-type and Gln143Asn MnSOD enzymes. Data are from (Hsieh *et al*., 1998).

## 2. Materials and Methods

### 2.1 Expression, purification, & crystallization of Gln143Asn MnSOD

The Gln143Asn plasmid was ordered from GenScript and codon optimized for expression in *Escherichia coli. (E. coli)*. Following transformation of *E. coli* BL21(DE3) cells with the plasmid, a small culture was incubated at 37 °C and 225 rpm until visible turbidity was reached in Terrific Broth (TB) supplemented with 30 µg/L of kanamycin. Next, 2 L of TB supplemented with 30 µg/L of kanamycin and 16 mL of 100% glycerol was inoculated with ∼5 mL of the small growth and incubated at 37 °C with vigorous shaking at 225 rpm until an OD_600_ of 0.8-1.0 was reached. A bolus of 8 mM MnCl_2_ and 1 mM isopropyl β-d-1-thiogalactopyranoside (IPTG) was then added, and the flasks were incubated at 37 °C and 140 rpm overnight. Induced cells were harvested by centrifugation using a Sorval Lynx 6000 with a Fiber Light F9-6x-1000 LEX rotor, operated at 4 °C and 8000 rpm for 40 minutes. The pellets were stored at –20 °C. For Gln143Asn MnSOD protein purification, pellets were thawed at room temperature and thoroughly resuspended in Lysis buffer (10 mM MnCl_2_ and 10 mM MOPS pH 7.8), followed by filtration and removal of the insoluble fraction. Next, the soluble fraction was lysed with an EmulsiFlex-C3 at a pressure equal to or greater than 20,000 psi for ∼5 sample passes. The lysate was centrifuged at 4 °C and 8,000 rpm for 40 minutes, and the supernatant was transferred to a hot water bath at 55 °C for 1 hour. Then, the solute was centrifuged again, and the supernatant (containing the soluble proteins) was diluted using 25 mM MES pH 5.5 at a 1:2 protein to buffer ratio. The diluted protein was then loaded onto a HiPrep 16×10 CMFF cation exchange column using an ÄKTA Pure HPLC (GE Healthcare). The sample was buffer exchanged into 10 mM MES at pH 5.5 on the column, followed by protein elution from the resin with 10 mM MES at pH 6.5. Purification was confirmed by the presence of a single band corresponding to the correct molecular weight of 23 kDa on a 4-12% SDS PAGE gel stained with Coomassie blue. Before crystallization, all pure fractions were pooled together, and buffer exchanged into 0.25 mM potassium phosphate (pH 7.8) using a 10 kDa concentrator (Millipore, Sigma Aldrich) and concentrated to ∼23 mg/ml (∼1 mM).

Twenty-four well hanging drop trays were set up with a 1:1 ratio of concentrated protein against reservoir solution containing 1.8 M potassium phosphate (pH 7.8) and incubated at room temperature. Gln143Asn MnSOD crystallized in the high-symmetry space group of P6_1_22, and the crystals grew to final sizes of ∼ 100 µm long in a week. The crystal trays were hand-carried by flight to the Stanford Synchrotron Radiation Lightsource (SSRL) for data collection.

### 2.2 X-ray crystallography data collection, processing, refinement, & validation

A single crystal of Gln143Asn MnSOD was exchanged into cryo-solution (2.5 M potassium phosphate, pH 7.8) at room temperature, manually plunged into liquid nitrogen, then mounted with a 100 µm-sized cryo-loop (MiTeGen) at SSRL beamline 14-1. The mounted crystal was annealed by manually blocking the cryo-stream for 5 seconds. For H_2_O_2_ soaking, a 30% stock solution of H_2_O_2_ (Millipore Sigma) was diluted 100x and pipetted into the drop of mother liquor containing the single Gln143Asn crystal, to reach a final H_2_O_2_ concentration of 0.3% *in-crystallo*. The total soaking time was ∼30 seconds at room temperature, followed by quick plunging into liquid nitrogen to cryo-trap the soaked H_2_O_2_ in the crystal, mounting, and annealing like the resting-state, resting state Gln143Asn MnSOD crystal. The process is schematized in Supplementary Information Fig. S1. X-ray crystallography data for both the resting-state and H_2_O_2_-treated Gln143Asnwere collected at 100 K, using a Dectris Pilatus 16M PAD detector and a controllable axial nitrogen cryo-stream (Oxford Cryosystems, Ltd.). Data collection parameters are summarized in Supplementary Information Table S1. Initial phases were obtained using the Phaser Molecular Replacement (Phaser-MR) package within the Phenix software suite with the wild-type MnSOD structure (PDB ID 7KLB) (Azadmanesh *et al*., 2021)serving as the phasing model. Model building was performed with COOT followed by iterative rounds of refinement cycles using Phenix_refine. All refinement strategies were based on maximum likelihood-based target functions and included the refinement of i) individual atomic coordinates (in both reciprocal and real space), and ii) individual isotropic atomic displacement parameters (ADPs) for all atoms, with automatic weight optimisation. Only riding hydrogens were added to the models (Liebschner, 2020) and appropriate metal coordination restraints were generated using idealised geometries, with the ReadySet tool. Gln143Asn, like the wild-type enzyme, crystallizes as a homodimer, with monomers A and B in the asymmetric unit, and crystallographic symmetry operations were applied to generate the physiological tetramer, using the PyMOL Molecular Graphics System (Schrodinger, LLC). Final model validations were performed using MolProbity. Coordinate error estimates, calculated with Phenix_refine, are reported Supplementary Information Table S1 as a measure of model precision. Coordinates and structure factors for the resting-state and H_2_O_2_-treated Gln143Asn MnSOD X-ray crystal structures have been deposited in the Protein Data Bank (PDB) under accession codes 9NR0 and 9NSJ, respectively.

### 2.3 Metal center specific electrostatic surface generation

To generate the solvent-excluded electrostatic surfaces of tetrameric MnSOD proteins that accurately reflect the electronic charge distribution, spin multiplicity, and redox potential of the active site Mn centers, we implemented a three-step protocol to account for electronic effects imparted by its primary coordination sphere (PCS) ligands (Van Stappen *et al*., 2022), schematized in Supplementary Information Fig. S2.

Step 1. Initial per-residue charges were assigned using the PDB2PQR (Dolinsky *et al*., 2007) web service, hosted at poissonboltzmann.org. This generated PQR files containing i) atomic partial charges of all residues, computed using the AMBER force field (Hornak *et al*., 2006) and ii) protonation states of ionizable residues, based on pKa values predicted by PROPKA 3.0 at pH 7.0 (Olsson *et al*., 2011).

Step 2. To determine the electronic properties of the catalytic Mn in specific redox and spin states, the protein structures were truncated to only retain the first-shell ligands coordinating the Mn ion, thereby preserving the key geometric and electronic features of the metal site. Density functional theory (DFT) calculations were performed on the isolated metal centers, using the high-spin sextet state for the divalent Mn ion in Gln143Asn enzyme (Azadmanesh *et al*., 2025), according to established protocols in (Neves *et al*., 2013, Azadmanesh *et al*., 2017). Single-atomic electrostatic potential (ESP) charges for the Mn ion and its direct ligands were derived using the Merz-Kollman scheme and subsequently refined using the restrained electrostatic potential (RESP) methodology. As a benchmark, we applied the same DFT protocol to the five-coordinate, high-spin quintet resting-state of the wild-type Mn^3+^SOD enzyme (PDB ID 5VF9) (Azadmanesh *et al*., 2017). All DFT-calculated ESP charges are given in Fig. 4C.

Step 3. The ESP-refined metal charges were then integrated with the remainder of the tetrameric protein’s PQR-derived charges to compute full electrostatic surface maps using the Adaptive Poisson-Boltzmann Solver (APBS) Electrostatics plugin within PyMOL, following the established protocol in (Azadmanesh *et al*., 2017).

### 2.4 Mn K-edge HERFD XANES spectroscopy data collection, processing, & analysis

A solution of 3 mM Gln143Asn MnSOD (∼70 mg ml^−1^) in 25 mM potassium phosphate (pH 7.8) was treated with 280 mM (1% *w/v*) H_2_O_2_ to isolate the H_2_O_2_-bound state of the Gln143Asn variant. Mn K-edge HERFD-XANES spectra (Proux *et al*., 2017) were recorded using a liquid He cryostat at 10 K, at beamline 15 -2 of SSRL. The incident energy was tuned to the first derivative of an internal Mn foil at 6539 eV. X-ray irradiation was carefully monitored so that two subsequent scans of the same spot did not have photoreduction differences, and different spots along samples were scanned. When appropriate, aluminium foil was inserted into the beam path to attenuate the incident flux. For HERFD-XANES measurements, a Johann-type hard X-ray spectrometer with six Mn analyser crystals were used with a liquid-nitrogen cooled Si (311) double crystal monochromator. Experimental spectra were processed, visualized, and plotted using the Larch software package. The post-edge regions were normalized, and pre-edge peak fittings were done using pseudo-Voigt functions. Pre-edge intensities were quantified as the integrated area under the fitted curves.

### 3.1 Multiple H_2_O_2_ binding sites were revealed beyond the Tyr34-His30 solvent gateway in Gln143Asn MnSOD

In this present study, we report two high-resolution X-ray crystal structures of the Gln143Asn MnSOD enzyme: (i) a 1.33 Å structure of H_2_O_2_-treated (PDB ID: 2NR0) and (ii) a 1.55 Å structure of the untreated resting-state (PDB ID: 2NSJ). Superposition of these two structures reveals that H_2_O_2_ soaking and cryotrapping triggers a striking accumulation of at least five distinct H_2_O_2_ molecules, either fully, or partially occupying the space between the solvent-exposed enzyme surface and the Tyr34-His30 solvent gate in Gln143Asn MnSOD (Fig. 2A). These H_2_O_2_ molecules in the bulk solvent are named L_b_ for Lig_bulk_. We compared L_b_ H_2_O_2_ (Fig. 2A,C, red sticks) to the corresponding water positions in the resting-state structure (Fig. 2D). Superposition shows that one or both oxygen atoms of the L_b_ H_2_O_2_ overlap with either fully or partially occupied water molecules in the untreated resting-state structure (Fig. 2D, blue spheres), indicating that the H_2_O_2_ molecules are hydrogen bonding to similar residues on the surface.

**Figure 2.**
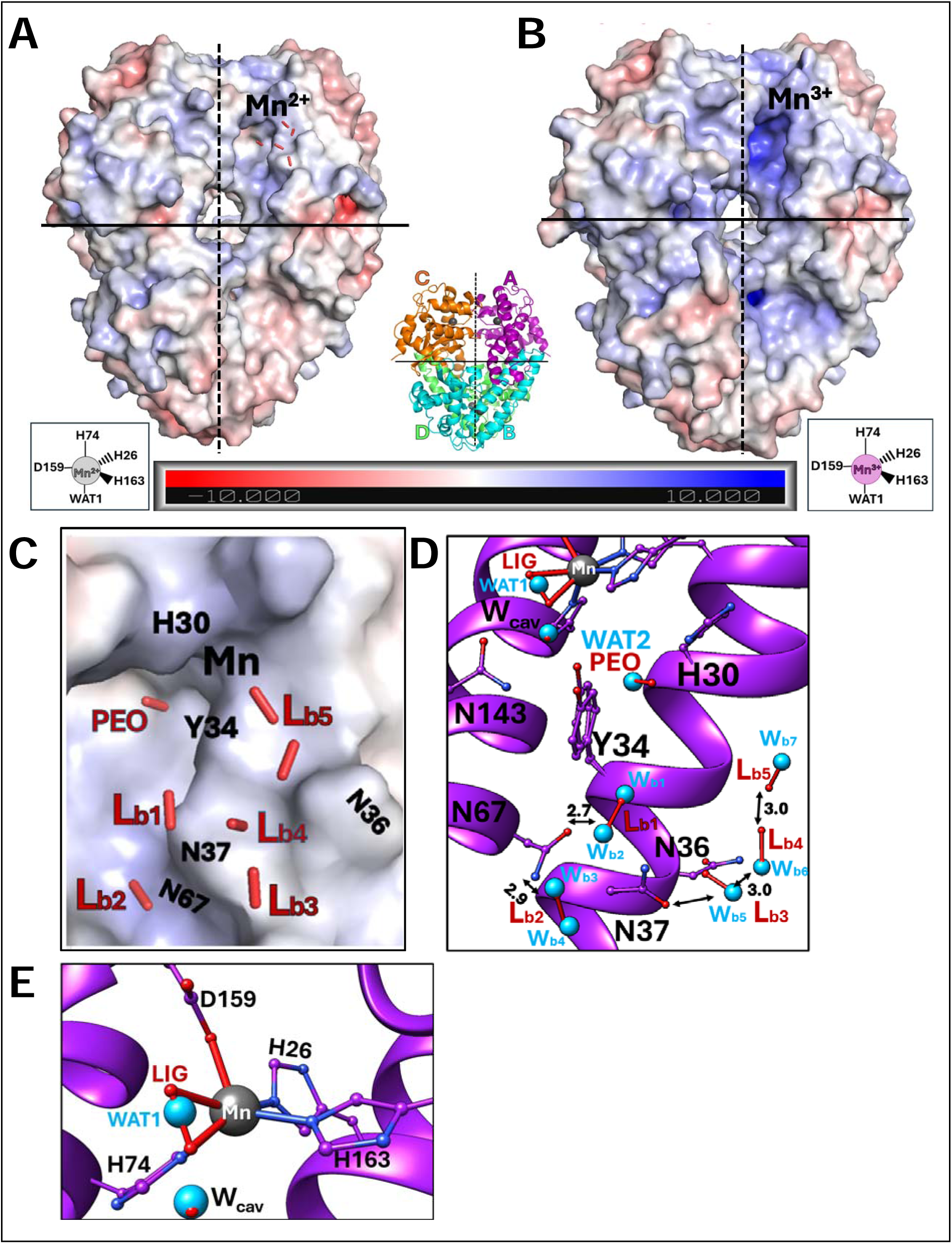
H_2_O_2_ binding. **(A-B)** Resting state Gln143Asn and wild-type MnSOD enzymes, overlaid on their respective electrostatic surfaces, with the location of the chain A active site Mn labelled. All bulk-exposed L_b_ peroxides trapped in the Gln143Asn enzyme are shown as red sticks. Dimeric and tetrameric interfaces are marked by dashed and solid black lines, respectively. Insets show the penta-valent Mn^2+^ (grey sphere) and Mn^3+^ (pink sphere) ions and their primary coordination sphere ligands in the Gln143Asn and wild-type enzymes, respectively. The middle inset shows the ribbon diagram of the tetrameric Gln143Asn MnSOD, with Mn ions shown as grey spheres. **(C)** Zoomed-in view of L_b_ H_2_O_2_ (red sticks), on top of the displayed electrostatic surface. **(D)** All modelled H_2_O_2_, active site (top), and the Tyr34-His30 solvent gate with the resting state water molecules (blue spheres). **(E)** Zoomed-in view of the active site of Gln143Asn, with bound LIG (red stick) peroxide in the treated structure, and the WAT1 solvent (blue sphere) and Wcav.

The L_b_ H_2_O_2_ molecules bind to the solvent-exposed surface of the Gln143Asn MnSOD tetramer (Fig. 2A,B). We note that the resting state electrostatic solvent accessible surface of Gln143Asn MnSOD is different than the surface of resting state wild-type MnSOD. The active site funnel of wild-type is much more positively charged than Gln143Asn. This is because the wild-type resting state is Mn^3+^, and the Gln143Asn resting state is Mn^2+^. The surface of wild-type MnSOD in the reduced state is practically identical to that of Gln143Asn. When wild-type MnSOD cycles its redox state during catalysis, its electrostatic surface cycles between the positive and more neutral surfaces. Gln143Asn will stay in the more neutral surface most of the time and this seems to help stabilize the H_2_O_2_ binding

To validate that the L_b_ electron densities observed in the H_2_O_2_-treated structure truly correspond to H_2_O_2_ molecules, rather than misassigned or mobile waters, we modeled fully occupied water (copied from the resting-state position) at each L_b_ site and studied the electron density after refinement. This analysis shows that each H_2_O_2_ site has an elongated 2Fo-Fc density. When modelled with a water molecule in trial refinements, the density is not satisfied, and residual Fo-Fc density remains (Supporting Information Fig. S3). This analysis confirms the identification of H_2_O_2_ binding sites. This many H_2_O_2_ molecules have never been observed in any previous MnSOD structure, and provide strong validation for our chosen experimental strategy, which leverages the slowed catalysis of the Gln143Asn variant to reveal all potential H_2_O_2_ binding sites in MnSOD.

In addition to the L_b_ H_2_O_2_ molecules on the solvent-exposed surface, we observe H_2_O_2_ binding at the metal-ligated LIG position, and at the PEO position at the mouth of the active site solvent-funnel gateway in between second-shell residues Tyr34 and His30. Overall, H_2_O_2_ treatment does not appear to significantly alter the active site architecture of Gln143Asn MnSOD, except for the partial displacement of the Mn-bound solvent (WAT1) by a partially occupied (∼35%) H_2_O_2_ molecule bound at the LIG position, opposite His26 (Fig. 2E). Superposition with all other previously published structures of H_2_O_2_-treated human MnSODs, i.e. wild-type (PDB ID 8VJ5) (Fig. 3A,B yellow), Trp161Phe (Fig. 3C cyan), and Tyr34Phe (Fig. 3C red), as well as *E.coli* wild-type MnSODs (PDB ID 3K9S) (Fig. 3A,B orange) shows that H_2_O_2_ binds the active site LIG position in the Gln143Asn variant (Fig. 3A-C, purple) in a configuration very similar to the previously observed nearly side-on binding modes in the human MnSOD variants, rather than end-on, as seen in the *E.coli* wild-type MnSOD.

**Figure 3.**
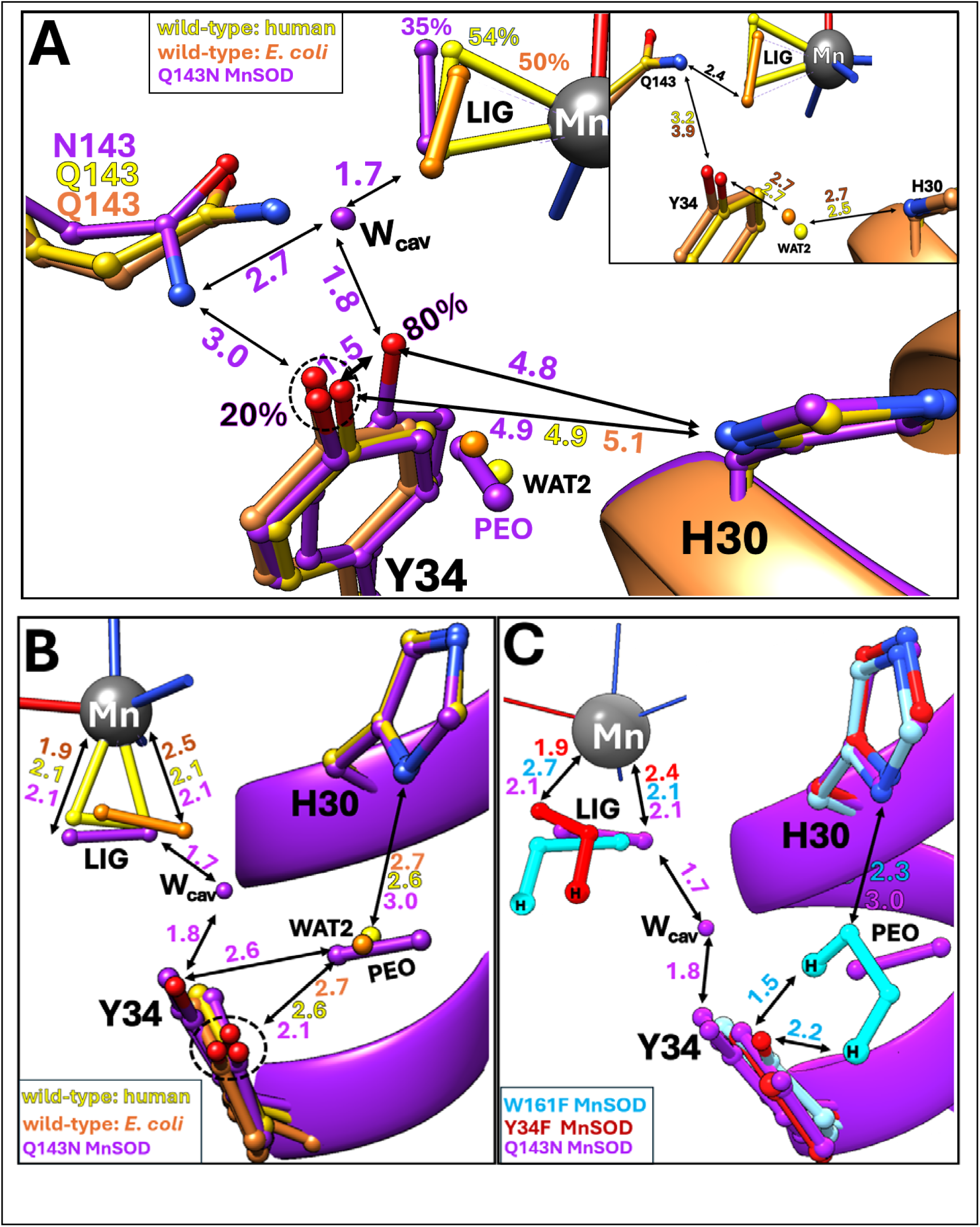
Comparison with known H_2_O_2_ soaked MnSOD in the PDB. Structural comparisons of H_2_O_2_ -treated human Gln143Asn MnSOD (purple) with (**A,B)** both human (yellow, PDB ID 8VJ5), and *E. coli* (orange, PDB ID 3K9S) wild-type enzymes, respectively, as well as with **(C)** the product inhibited MnSOD variants, Trp161Phe (cyan, neutron PDB ID 8VHW) and Tyr34Phe (red, neutron PDB ID 9BVY). We observe H_2_O_2_ binding in the pre-established LIG and PEO binding sites, with subtle differences, potentially owing to the Gln143 mutation-induced cavity. Notably, Tyr34 samples a shifted conformation (∼80% occupancy) in this cavity, drawing closer to both the LIG H_2_O_2_ and His30. This conformational flexibility is not typically observed in wild-type or other variants lacking this mutation-induced cavity, between the metal primary sphere and the second-shell solvent gateway.

**Figure 4.**
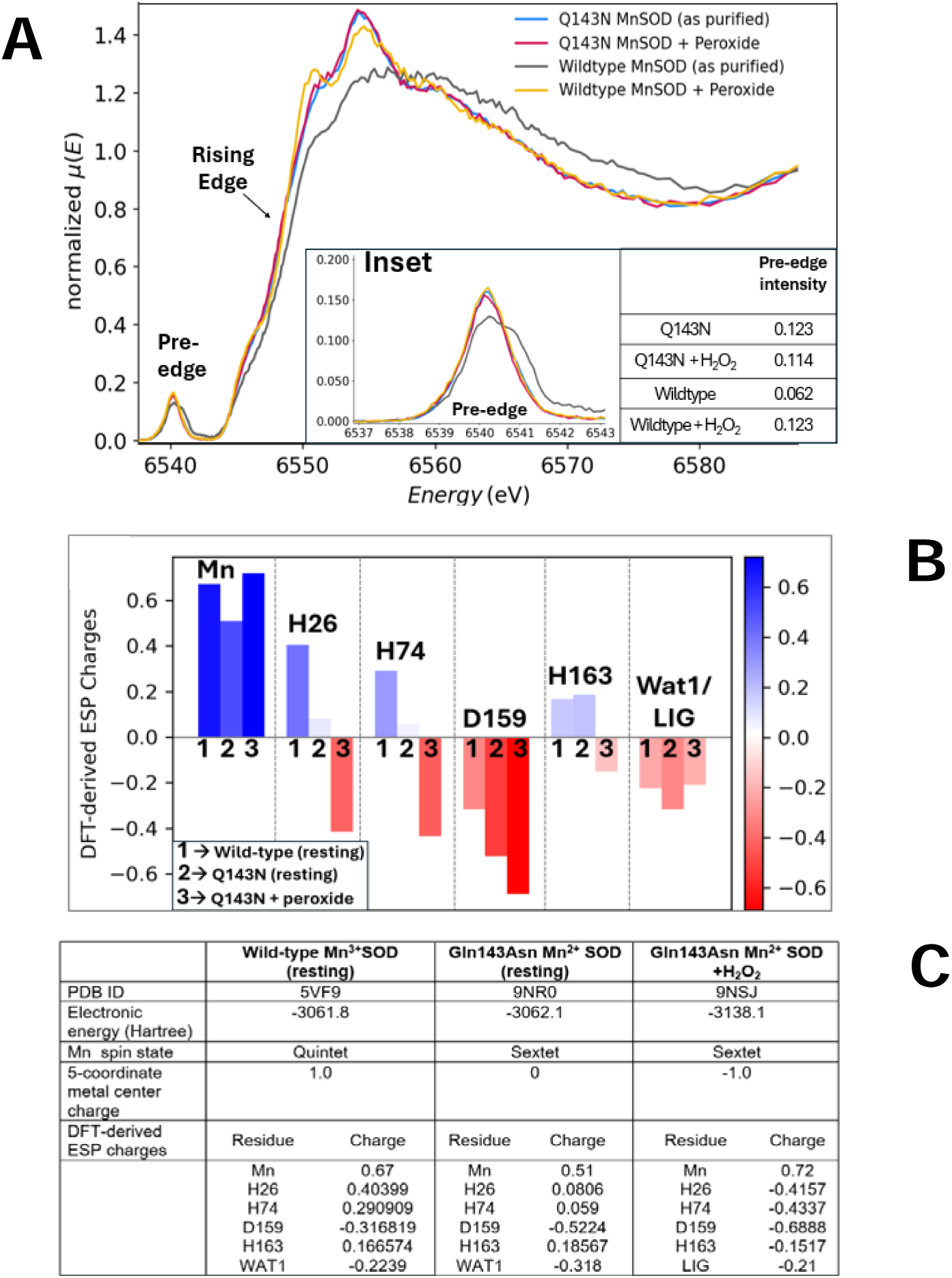
HERFD XANES spectroscopy **(A)** Normalized Mn K-edge HERFD-XANES spectra of wild-type and Gln143Asn MnSOD enzymes, both in resting and H_2_O_2_-soaked forms. **Inset** shows a magnified view of the pre-edge region (6537–6543 eV), along with their individual pre-edge intensities. **(B and C)** Mn redox state and/or H_2_O_2_ treatment affects the individual DFT-derived ESP charges of the Mn primary coordination sphere residues.

Additionally, we observe a 57% occupied H_2_O_2_ near the PEO binding site, which has so far only been observed in the H_2_O_2_-soaked neutron structure of Trp161Phe MnSOD (PDB ID 8VHW) (Azadmanesh *et al*., 2024), at full occupancy (Fig. 3C cyan). Superposition of the two H_2_O_2_-bound MnSOD structures reveals that H_2_O_2_ binds at the PEO site of the Gln143Asn variant (Fig. 3C, purple) at a slightly different configuration. However, we do not know what causes this change in H_2_O_2_ binding at the PEO site of Gln143Asn MnSOD.

### 3.2 Peroxide binding subtly alters the active site electrostatics Gln143Asn MnSOD without changing the redox or coordination states of the catalytic Mn

To monitor changes in the oxidation state and coordination geometry of the catalytic Mn in the Gln143Asn variant upon H_2_O_2_ treatment, we compared the Mn K-edge High-Energy-Resonance Fluorescence-Detected X-ray Absorption Near-Edge Spectra (HERFD-XANES) of the purified enzyme, before and after H_2_O_2_ treatment, following protocols established in (Azadmanesh *et al*., 2025, Azadmanesh *et al*., 2024). We observe no detectable shift in either rising edge or pre-edge features (Fig. 4A blue vs red), indicating no alteration in the redox state or coordination geometry of the catalytic Mn after H_2_O_2_ treatment in Gln143Asn MnSOD. Notably, the spectral features of both Gln143Asn samples closely resemble those of the H_2_O_2_-treated wild-type MnSOD (Fig. 4A yellow), which has been previously established to contain the reduced Mn^2+^ ion in its active site (Azadmanesh *et al*., 2024). The pre-edge intensities of both Gln143Asn samples are comparable to each other and the H_2_O_2_-treated wild-type enzyme, indicating highly similar metal centers across all three conditions. In contrast, the spectrum of the resting-state wild-type MnSOD (Fig. 4A, black) shows a clear shift of both rising and pre-edge features toward higher energies, consistent with the established presence of an oxidized Mn^3+^ metal in the resting wild-type MnSOD.

DFT calculations predict no significant change in the charge or spin state of the catalytic Mn^2+^ in Gln143Asn enzyme, upon H_2_O_2_ treatment. However, certain trends emerge (Fig. 4BC): (i) Charges of both His26, and His74 switch signs, going from positive to negative (and closer to the wild-type resting-state values), (ii) Asp159 is less negatively charged in the resting-state of Gln143Asn, compared to wild-type, but becomes more negative, with H_2_O_2_ addition, and (iii) His163 flips from negative to positive charge when comparing the wild-type to the variant, indicating potential protonation state change. However, these conclusions remain to be experimentally validated. Moreover, it is important to note that our DFT models treat LIG as a di-oxygen species rather than H_2_O_2_, to account for the caveat that X-ray electron density maps cannot pinpoint proton positions accurately. This is particularly relevant considering recent neutron diffraction studies of H_2_O_2_-soaked Trp161Phe (Fig. 3C cyan), and Tyr34Phe (Fig. 3C red) Mn^2+^SODs, both of which stabilize a singly protonated di-oxygen species, hydroperoxyl anion (^−^OOH), following H2O2-induced metal reduction (Azadmanesh *et al*., 2025, Azadmanesh *et al*., 2024).

Overall, we conclude that H_2_O_2_ treatment does not alter the redox state or coordination geometry of the catalytic Mn in the Gln143Asn enzyme, unlike all other H_2_O_2_-treated MnSOD variants characterized previously (Azadmanesh *et al*., 2025, Azadmanesh *et al*., 2024). This further demonstrates how Gln143Asn is perpetually frozen in the reduced state without the necessary proton transfer events. This loss of redox flexibility at the expense of catalytic turnover distinguishes Gln143Asn from all other spectrally characterized human MnSOD variants to date.

## 4. Conclusions

The high-resolution crystal structure of H2O2-treated human Gln143Asn MnSOD reported herein reveals previously unobserved, L_b_ H_2_O_2_ binding sites on the solvent-exposed surface in the bulk solvent, in addition to the canonical LIG and PEO positions (Fig. 2&3). This contrasts with all previously published H_2_O_2_-soaked MnSOD structures, which have captured H_2_O_2_ binding only within the first or second coordination shells. Although the fact that product release occurs too rapidly in the wild-type enzyme offers a plausible explanation as to why we do not catch transient H_2_O_2_ beyond the active site gateway on the solvent-exposed side, their absence even in severely product-inhibited MnSOD variants, such as Trp161Phe and Tyr34Phe, suggests that the principal kinetic bottleneck for product release from MnSOD is H_2_O_2_ diffusion across the Tyr34-His30 solvent gate. It appears that once beyond this point, H_2_O_2_ escapes rapidly into bulk solvent, evading experimental detection methods. To overcome this gap, we leveraged the slow kinetics of the catalytically impaired Gln143Asn MnSOD to successfully stabilize several L_b_ H_2_O_2_ beyond the active site solvent funnel, thus depicting a more extensive MnSOD H_2_O_2_-interaction landscape than previously available (Fig. 2). The redox cycling of the active site metal in Gln143Asn MnSOD is impaired, this causes the electrostatic surface to stay in a more neutral condition and the H_2_O_2_ are fairly immobilized (Supporting Information Fig. S3). These solvent-exposed L_b_ H_2_O_2_ likely represent a native ensemble of product molecules diffusing across the active site solvent channel, extending from the gateway at the second shell, to the bulk solvent-exposed surface. This phenomenon has never been structurally visualized before in any H_2_O_2_-soaked MnSOD enzyme.

Tyr34 in Gln143Asn adopts a dominantly shifted conformation (∼80% occupancy), even in the absence of H_2_O_2_ soaking (Fig. 2D, 3). This conformation is stabilized by an additional solvent molecule position in the mutation-induced cavity, namely, W_cav_, which in turn orients toward the LIG position-bound species, creating a tight interaction network that could stabilize bound H_2_O_2_ at the LIG site. This network is not observed in the wild-type or other MnSOD variants, due to steric hindrance from Gln . This water molecule might be able to donate a proton to WAT1, and this may be why Q143N retains some activity. The alternate Tyr34 conformation and the presence of the W_cav_ solvent introduce new hydrogen bonds within the active site of Gln143Asn MnSOD, which are not observed in the wild-type enzyme (Supporting Information Fig. S6). These may contribute to the slightly higher thermal stability of Gln143Asn MnSOD (∼1.8 °C) previously reported (Hsieh *et al*., 1998).

HERFD-XANES and DFT analyses show no detectable change in the oxidation state, coordination number, or spin state of the catalytic Mn²lJ center of Gln143Asn upon HlJOlJ treatment (Fig. 3A). These results confirm that this MnSOD variant remains stuck in a reduced, catalytically inert state, unable to drive the PCET-mediated MnSOD catalytic cycle. By decoupling proton transfer from catalysis, the Gln143Asn mutation stabilizes transient H_2_O_2_-bound states and opens a rare structural window into otherwise elusive steps of MnSOD catalysis. This work underscores the value of catalytically impaired mutants not as dysfunctional models, but as powerful structural probes for revealing elusive mechanistic events in metalloenzymes.

## Acknowledgements

This research was funded by the NIGMS grant R01-GM145647. The UNMC Structural Biology Core Facility received support from the Fred and Pamela Buffett NCI Cancer Center Support Grant (P30CA036727). The use of the Stanford Synchrotron Radiation Lightsource (SSRL) at SLAC National Accelerator Laboratory, is supported by the US Department of Energy (DOE), Office of Science, Office of Basic Energy Sciences under Contract No. DE-AC02-76SF00515. The SSRL Structural Molecular Biology Program is funded by the DOE Office of Biological and Environmental Research and the NIH National Institute of General Medical Sciences (P30GM133894). The views expressed in this publication are those of the authors and do not necessarily reflect the official positions

## Supporting information

**Table S1.**
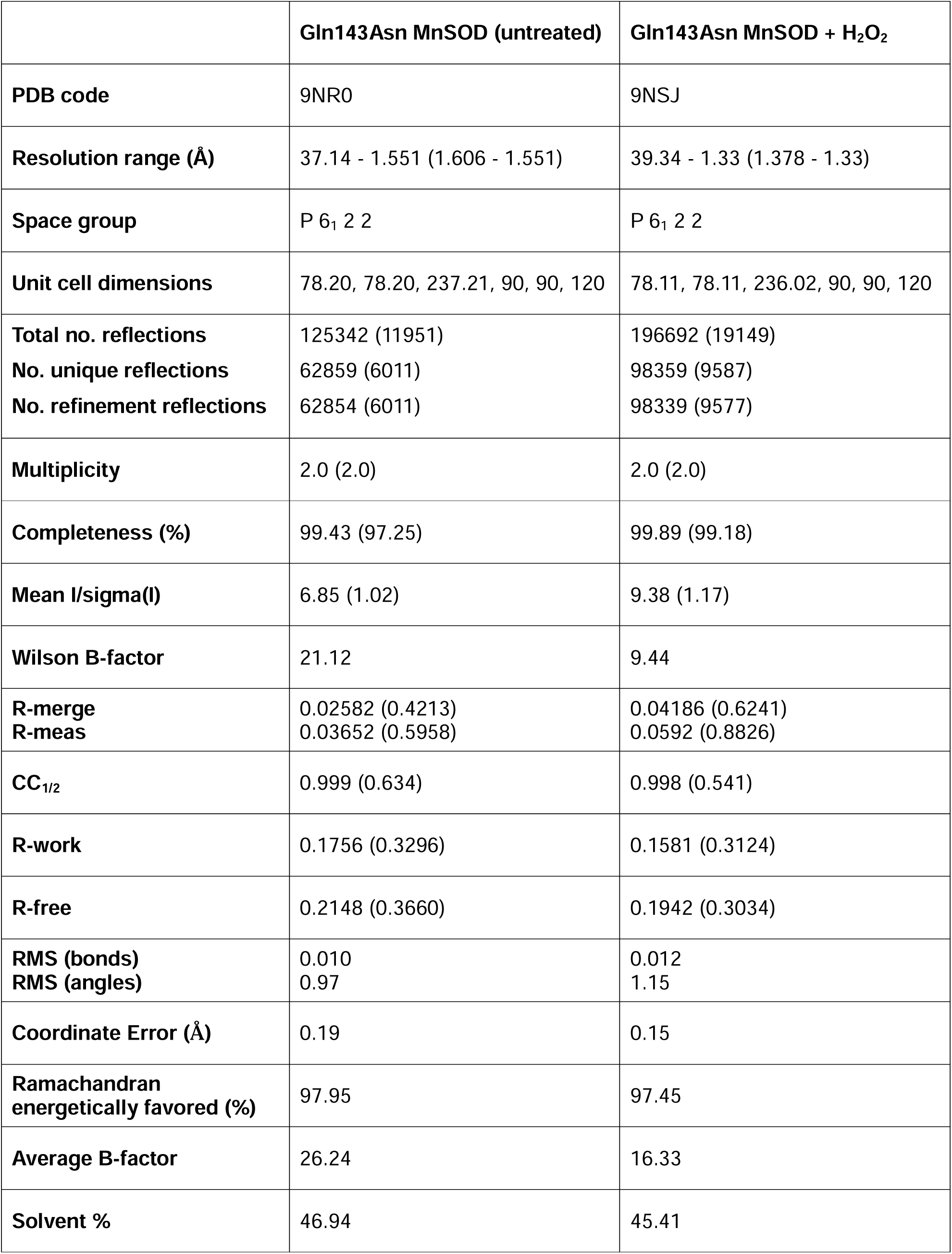
X-ray data collection and refinement statistics.

**Table S2.**
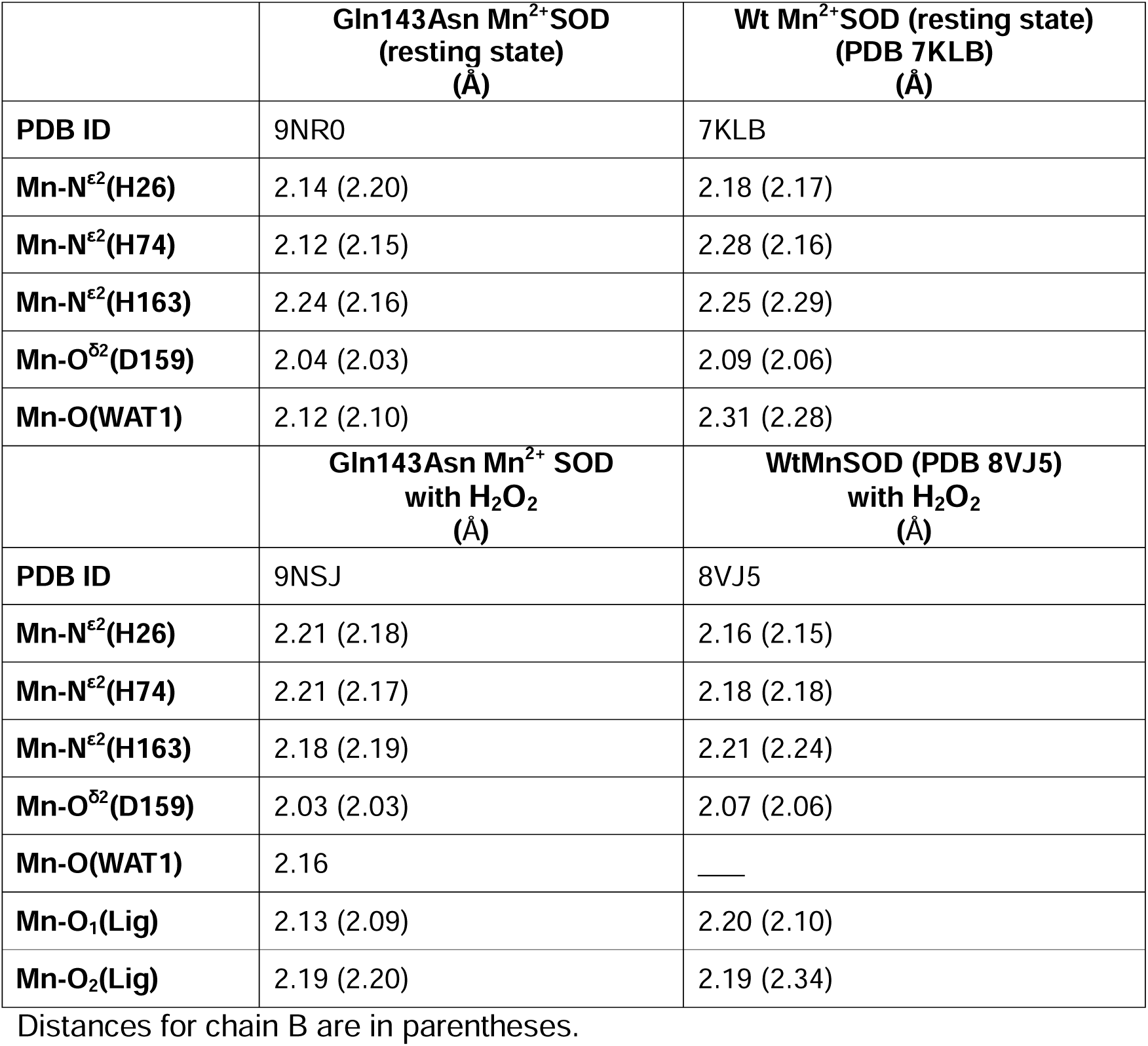
Active site Mn bond lengths in resting-state and H_2_O_2_-bound Gln143Asn Mn^2+^ SOD and wildtype Mn^3+^ SOD.

**Table S3.** Active site water (W_cav_) coordination in resting-state and H_2_O_2_-treated Gln143Asn MnSOD.

|  | **Gln143Asn MnSOD**<br>**untreated**<br>**(Å)** | **Gln143Asn MnSOD**<br>**with $\text{H}_2\text{O}_2$**<br>**(Å)** |
| --- | --- | --- |
| **PDB ID** | 9NR0 | 9NSJ |
| **$\text{O}^{\text{H}}(\text{Y34}) - \text{O}(W_{\text{cav}})$** | 2.01 (2.10) | 1.83 (1.94) |
| **$\text{N}^{\delta 2}(\text{N143}) - \text{O}(W_{\text{cav}})$** | 2.79 (2.47) | 2.66 (2.52) |
| **$\text{O}(\text{WAT1}) - \text{O}(W_{\text{cav}})$** | 2.24 (2.35) | 2.41 |

Distances for chain B are in parentheses.

**Figure S1.**
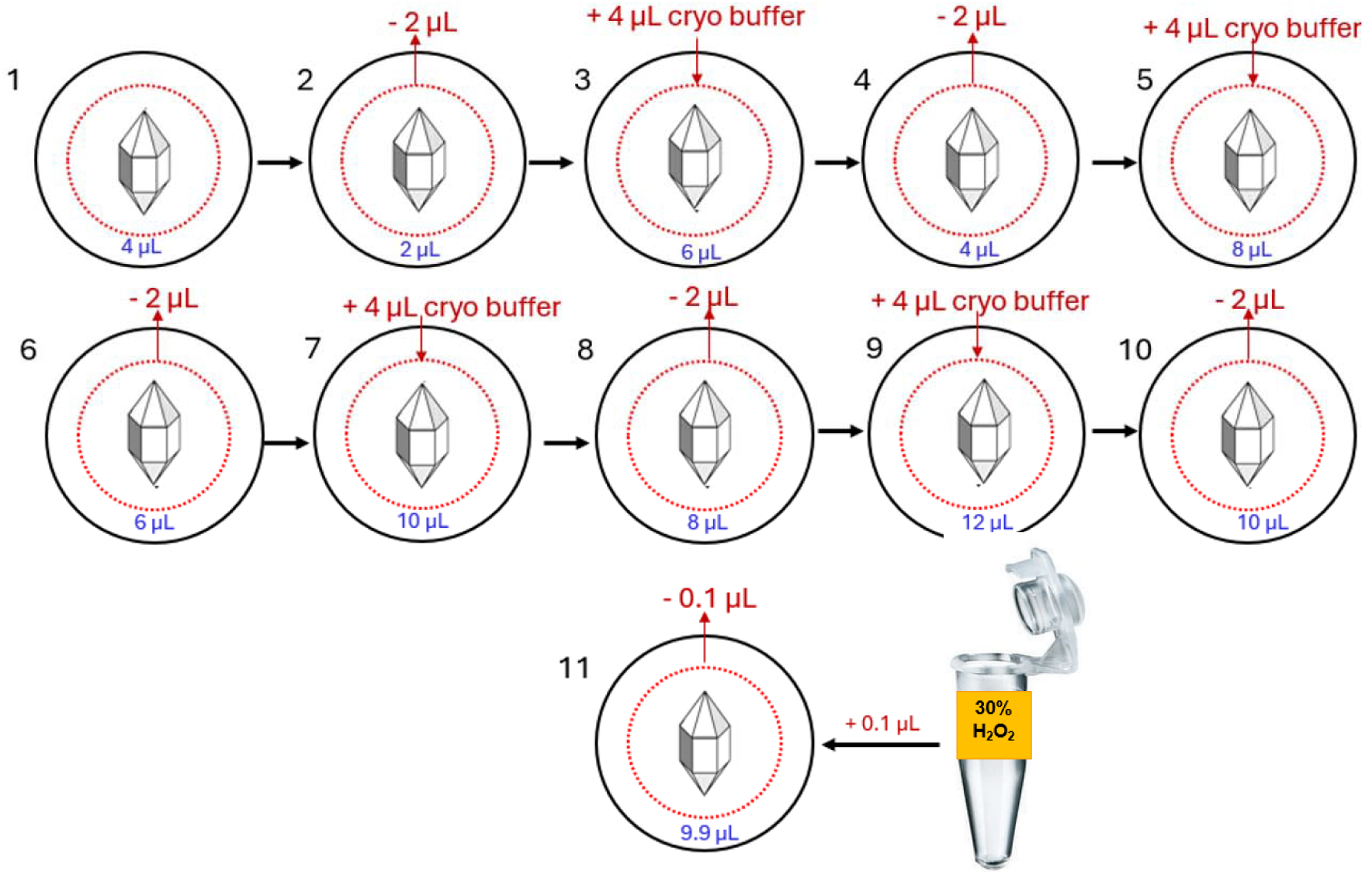
**Cryoprotection protocol for Gln143Asn MnSOD crystals**: The crystallization drop is shown as a red dashed circle with the final drop volume labeled in blue. The direction of red arrows shows if liquid is pipetted away from the crystal drop, or if cryo-buffer is pipetted into the drop. Only for the H_2_O_2_-treated Gln143Asn MnSOD dataset, 0.1µL of a 30% stock of H_2_O_2_ (Sigma Aldrich) was added to a total drop volume of 10 µL to reach a final H_2_O_2_ concentration of 0.3% in the single crystal (step 11) and allowed to soak for ∼30 seconds at room temperature. This was followed by plunging into liquid nitrogen and mounting in the cryostream at SSRL 14-1.

**Figure S2.**
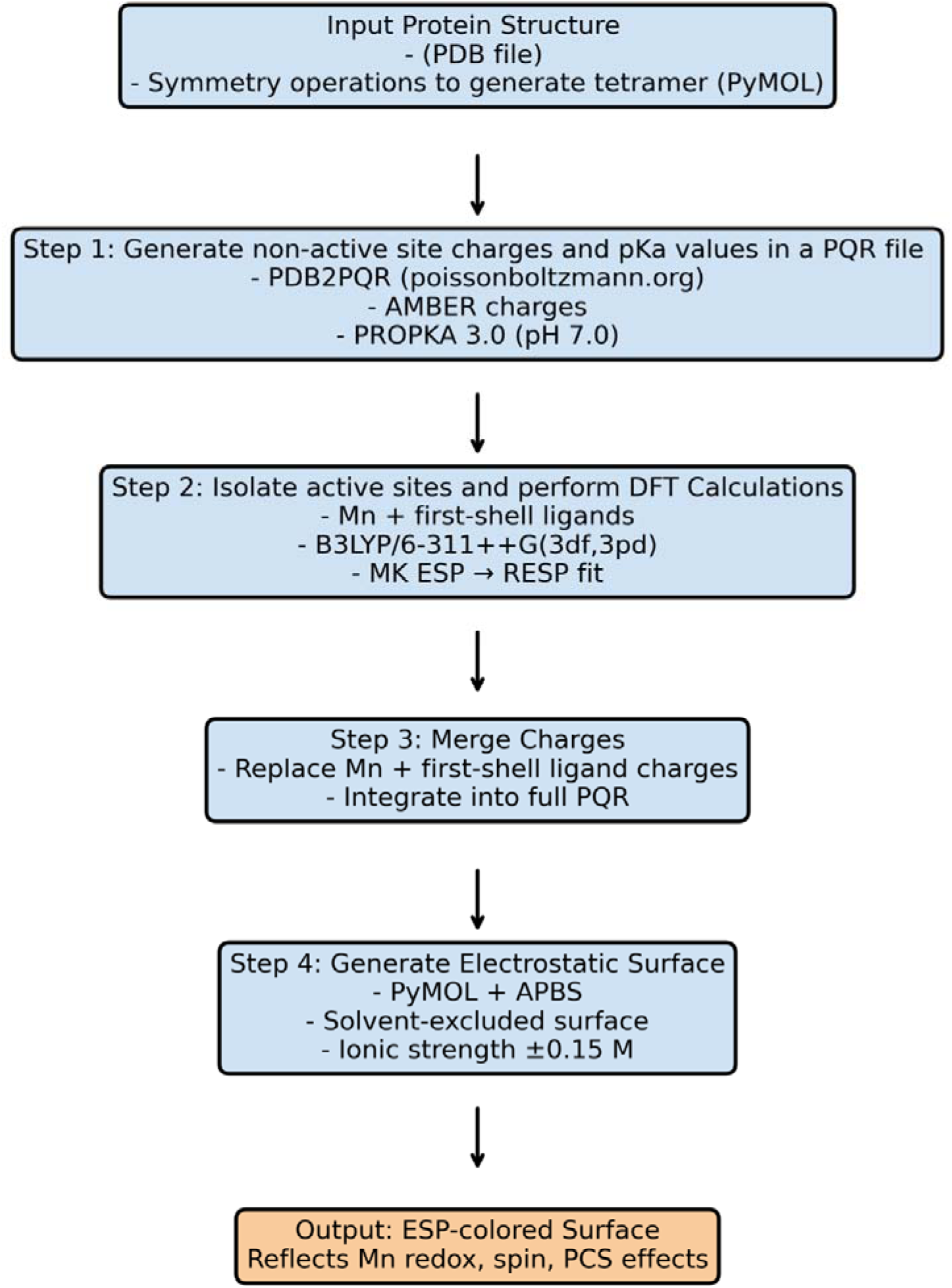
Workflow for generating metal-centered electrostatic potential maps.

**Figure S3.**
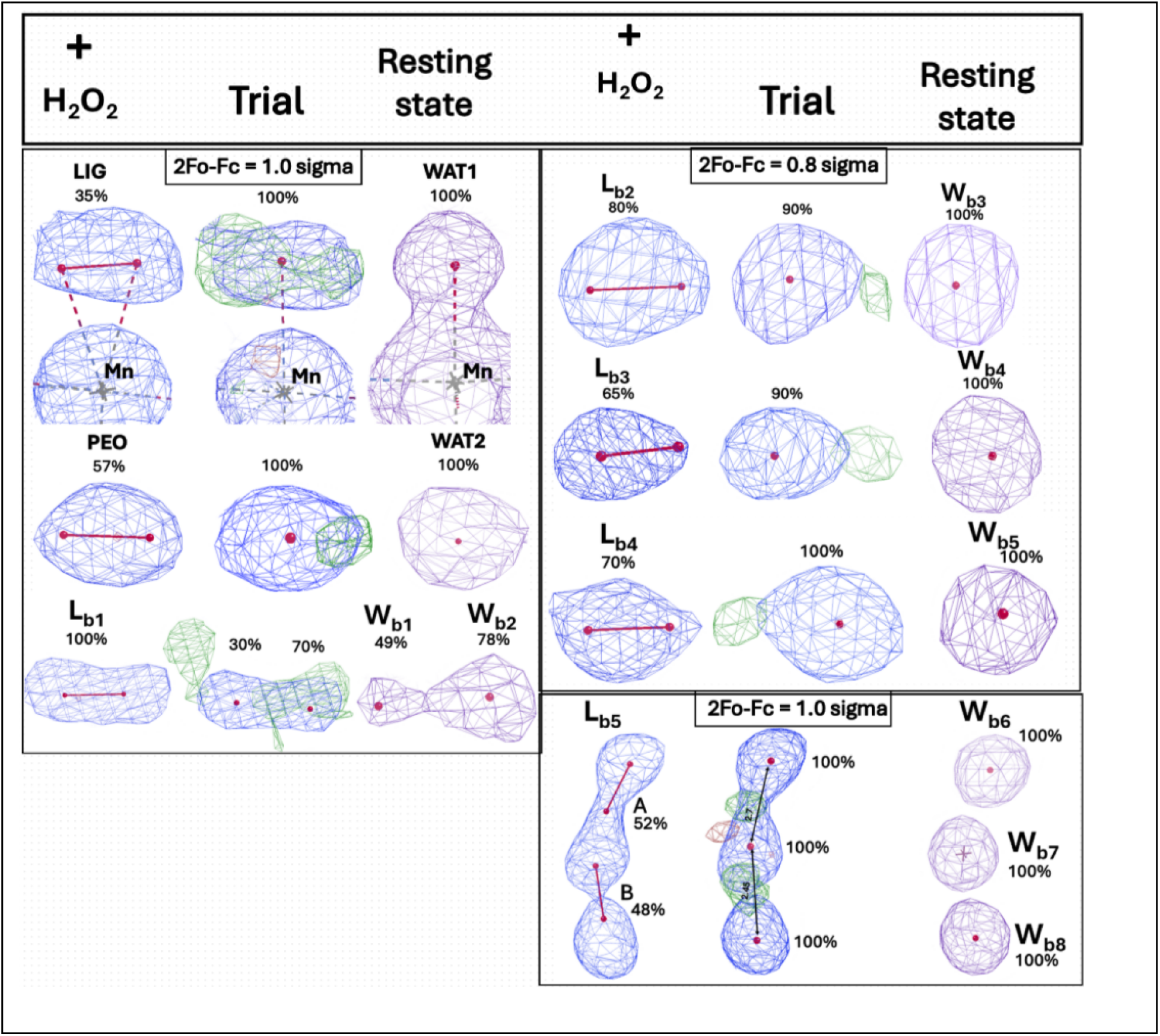
Solvent-exposed bulk peroxide (L_b_) modeling trials in Gln143Asn MnSOD. To rigorously validate the H_2_O_2_ models, L_b_ electron densities, a refinement trial was performed where fully occupied water molecule positions from the nontreated, resting-state Gln143Asn MnSOD structure were placed at each L_b_ sites. 2Fo-Fc electron density maps for modeled L_b_ H_2_O_2_ in the H_2_O_2_-treated structure are shown in blue mesh, and corresponding waters in the resting state structure are a purple mesh. The 2Fo-Fc density is elongated for the L_b_ sites, and the water are spherical. The Fo-Fc difference densities (green mesh, contoured to 3.0 sigma) in the middle trial panels indicate the need to model two oxygen atoms, rather than one.

**Figure S4.**
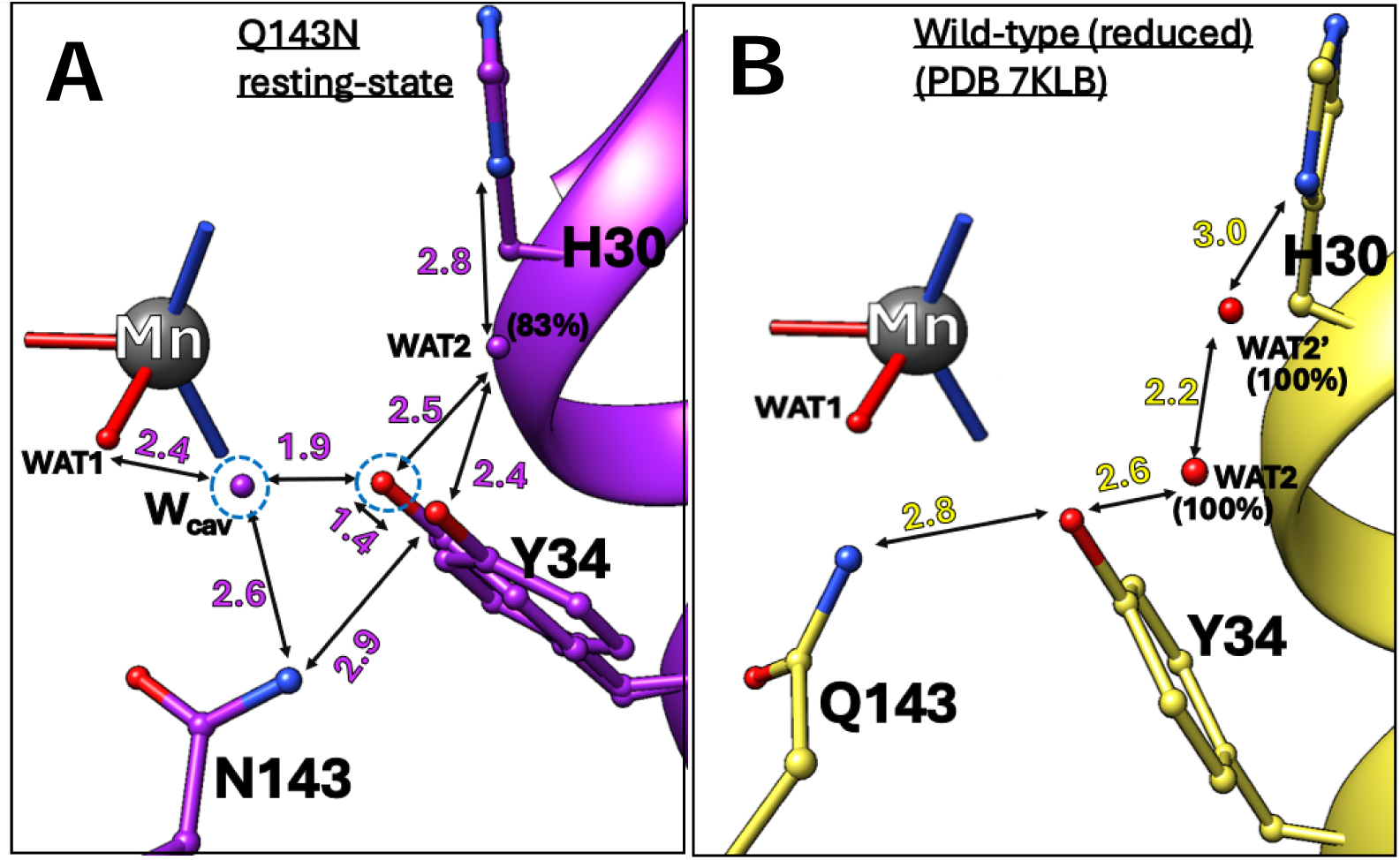
The Gln143 to Asn mutation creates a cavity adjacent to the active site which accommodates Tyr34 conformational flexibility in Gln143Asn MnSOD. (A) In the Gln143Asn MnSOD structure, the mutation-induced cavity accommodates a second conformation of Tyr34 (80% occupancy, circled in blue) and an additional water molecule, W_CAV_ (also circled in blue), resulting in new hydrogen bonding interactions. WAT2, located between Tyr34 and His30, is shown as a purple sphere. (B) The same view in reduced wild-type MnSOD (PDB ID: 7KLB) shows no evidence of a second Tyr34 conformation or W_CAV_ and has significantly fewer hydrogen bonds overall. These differences may contribute to the observed ∼1.8ℒ°C increase in thermal stability of Gln143Asn MnSOD relative to the wild-type enzyme.

**Figure S5.**
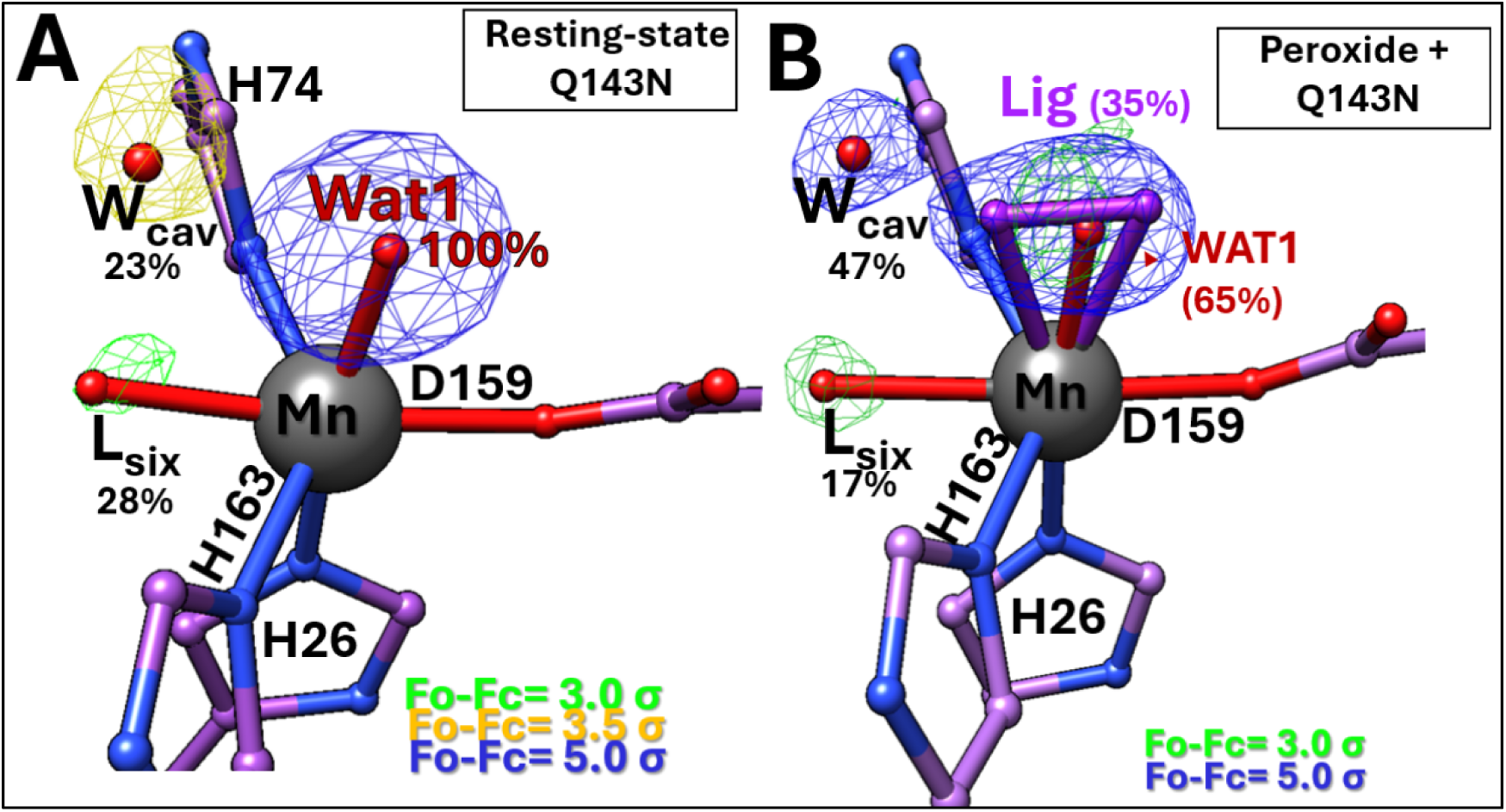
Gln143Asn MnSOD supports a partially hexavalent Mn coordination environment. **(A)** and **(B)** show the resting-state and H_2_O_2_ -treated active sites of Gln143Asn MnSOD respectively. display significant difference electron densities consistent with the presence of partially occupied sixth ligand (named L_six_), likely a hydroxide ion (OH^-^), opposite Asp159, observed at 23% occupancy in the resting state, and 17% following H_2_O_2_ exposure. This sixth ligand has previously been reported, only at full occupancy in chain A of the reduced human wild-type Mn^2+^SOD (PDB 7KKW). Difference densities for the modelled W_cav_ and L_six_ in both structures of the Gln143Asn are represented as mesh, coloured according to their individual contour levels. Presence of this partial sixth coordination partner induces a partial distortion of the Mn^2+^ geometry toward octahedral, diverging from the canonical trigonal bipyramidal configuration characteristic of the penta-coordinate active site

